# Evolutionary history of an Alpine archaeognath (*Machilis pallida*) – insights from different variant types

**DOI:** 10.1101/2021.09.17.460766

**Authors:** Marlene Haider, Martin P. Schilling, Markus H. Moest, Florian M. Steiner, Birgit C. Schlick-Steiner, Wolfgang Arthofer

## Abstract

Reconstruction of species histories is a central aspect of evolutionary biology. Patterns of genetic variation within and among populations can be leveraged to elucidate evolutionary processes and demographic histories. However, interpreting genetic signatures and unraveling the contributing processes can be challenging, in particular for non-model organisms with complex reproductive modes and genome organization. One way forward is the combined consideration of patterns revealed by different molecular markers (nuclear vs. mitochondrial) and types of variants (common vs. rare) that differ in their age, mode and rate of evolution. Here, we applied this approach to *Machilis pallida* (Archaeognatha), an Alpine jumping bristletail considered parthenogenetic and triploid. We generated de-novo transcriptome and mitochondrial assemblies to obtain high-density data to investigate patterns of mitochondrial and common and rare nuclear variation in 17 *M. pallida* individuals sampled across the Alps from all known populations. We find that the different variant types capture distinct aspects of the evolutionary history and discuss the observed patterns in the context of parthenogenesis, polyploidy and survival during glaciation. This study highlights the potential of different variant types to unravel complex evolutionary scenarios and the suitability of *M. pallida* and the genus *Machilis* as a study system for the evolution of sexual strategies and polyploidization during environmental change. We also emphasize the need for further research which will be stimulated and facilitated by these newly generated resources and insights.

## 1. Introduction

A central aspect of evolutionary biology is the reconstruction of species and population histories using fossil, morphological and genetic data. Patterns of genetic variation within and among populations are shaped by evolutionary and demographic processes and carry information which can be leveraged to infer the evolutionary history of natural populations and species. These signatures can be useful to reconstruct population structure, changes in population size, dispersal events and migration, as well as admixture and selection. Moreover, changes in reproductive mode (e.g. switches from sexual to parthenogenetic reproduction) or genome organization (e.g. polyploidization) leave detectable traces in the genetic makeup of populations and species. However, interpreting these signals and correctly assigning them to the aforementioned processes can be challenging. Such studies often require high-quality data as well as extensive modelling and simulations as well as accurate estimates of population parameters (e.g. mutation and recombination rate and generation time), which are usually only available for a few intensively studied organisms. Despite excellent theoretical work covering the population genetics of organisms that deviate from sexual reproductive strategies (e.g. Muller, 1964; Hill and Robertson, 1966; Felsenstein, 1974; Birky, 1996; Charlesworth and Charlesworth, 1997; Gerrish and Lenski, 1998; Barton and Charlesworth, 1998) and diploidy (e.g. White, 1973; Maynard Smith, 1978; Burch and Jung, 1993; Dufresne and Hebert, 1994), disentangling the forces that shape genetic diversity remains difficult (Ellegren and Galtier, 2016; Tellier, 2019).

An alternative approach, which is applicable to non-model organisms as well, is the combined consideration of patterns revealed by different molecular markers and/or types of variants that differ in their age, mode, and rate of evolution. A common practice is the comparison of patterns reflected by nuclear and cytoplasmic molecular markers found in mitochondria (mtDNA) in animals, mitochondria and chloroplasts (ptDNA) in plants, various organelles in fungi and protists, and even intracellular bacteria. Those markers differ from nuclear markers in their effective population size (Ne), recombination as well as mutation rate, and mode of inheritance. For example, mitochondrial and chloroplast markers are subject to stronger drift due to their lower Ne and have therefore frequently been used to investigate recent colonization events and changes in the connectivity among populations (Harrison, 1989; Schönswetter et al., 2005). In addition, incongruence of phylogenetic signals found in nuclear and mtDNA/ptDNA can inform about gene flow events (Avise et al., 1987; Funk and Omland, 2003; Baldo et al., 2011).

When high-density markers are available, a different approach can be applied that makes use of the fact that patterns of common and rare variants are affected differently by population genetic and demographic processes. On the one hand, common variants are expected to be shared among many individuals and populations and therefore likely reflect broader geographic patterns and older events. On the other hand, many rare variants likely reflect recent mutations that have not yet spread or could be older variants in a migration-drift ‘quasi-equilibrium’ (Slatkin, 1985; Slatkin and Takahata, 1985; Barton and Slatkin, 1986)(reviewed in Gompert et al., 2014). Regardless of the underlying processes, however, these variants are expected to be spatially restricted under low dispersal and gene flow (Barton and Slatkin, 1986)(discussed in Gompert et al., 2014). In fact, several studies found that rare variants reflect more recent and geographically localized processes in humans (Li et al., 2010; Mathieson and McVean, 2012; Nelson et al., 2012; Mathieson and McVean, 2014). Moreover, this pattern seems to be congruent in Lycaeides butterflies (Gompert et al., 2014), *Arabidopsis* (Memon et al., 2016), and *Boechera* rockcress (Schilling et al., in prep.) as well.

The aforementioned approaches utilizing rare and common nuclear variants as well as mitochondrial markers can be combined to leverage the differences in resolution and sensitivity to population genetics processes of these three types of variants. We use this highly versatile framework to gain a better understanding of the complex evolutionary history of an Alpine apterygote insect and extend it to reassess previously proposed scenarios of switches in reproductive mode and ploidy (Wachter et al., 2012).

The jumping bristletail genus *Machilis* (Archaeognatha or Microcoryphia) comprises 94 described species and with 55 species, the European Alps harbor the highest species diversity (de Jong et al., 2014; Dejaco et al., 2016). Due to their ancestral winglessness and multiple shared plesiomorphies with other insect orders they have often been described as ‘ancestral’ or ‘primitive’(Sturm and Machida, 2001). Given the lack of wings and the high degree of endemism in the genus, these bristletails are assumed to be slow dispersers (Sturm and Machida, 2001; Dejaco et al., 2016), however, modes of dispersal as well as the potential for dispersal are still unknown. To date, exclusively females have been found for several species and populations despite intensive sampling campaigns (Janetschek, 1956; Palissa, 1964; Sturm and Machida, 2001; Wachter et al., 2012; Rinnhofer et al., 2012; Dejaco et al., 2012, 2016), strongly suggesting the occurrence of parthenogenetic reproduction in the genus. Moreover, karyotyping and flow cytometry results indicate various instances of polyploidy (Gassner et al., 2014). Although polyploidization is strongly correlated with parthenogenetic reproduction in animals, this is not a general rule (Otto and Whitton, 2000), and there is also no such strict association in *Machilis*. In the species studied so far, sexuals were found to be diploid whereas asexuals were classified as either diploid or triploid (Gassner et al., 2014).

The high number of *Machilis* endemics and their distribution in and around the European Alps has sparked interest in their distribution and survival during the ice ages. Identifying and characterizing the different refugia during the last glacial maximum (LGM; 18,000 years before present)(van Husen, 1997) enables a better understanding of current species distributions and levels of diversity, both in terms of species numbers and genetic variation (Knowles, 2000; Schönswetter et al., 2002, 2005; Stehlik, 2003; Holderegger and Thiel-Egenter, 2009; Westergaard et al., 2011; Schneeweiss and Schönswetter, 2011). Such information also increases our knowledge on how species can react to climate change and which biological features are associated with successful survival in a changing environment. One open debate in this context revolves around the question of whether and how frequently arctic and high alpine species survived glaciation on nunataks (isolated mountain top areas protruding above the ice sheet) versus in peripheral areas (Schoönswetter et al., 2005; Schneeweiss and Schönswetter, 2011). While several studies in plants suggest nunatak survival (Stehlik et al., 2002; Abbott and Brochmann, 2003; Bettin et al., 2007; Parisod and Besnard, 2007; Westergaard et al., 2011), there is little evidence in animals, with the exception of *Trechus* ground beetles from peripheral nunataks in the Orobian Alps (Lohse et al., 2011).

One case in animals, for which central nunatak survival has been suggested (Wachter et al. 2012) is *Machilis pallida* (Janetschek, 1949), an endemic bristletail in the Eastern Alps, residing exclusively on carbonate rock scree above 2000 m above sea level (a.s.l.) (Dejaco et al., 2012; Rinnhofer et al., 2012; Wachter et al., 2012). The species is considered to be parthenogenetic (Rinnhofer et al., 2012; Wachter et al., 2012) and triploid (Gassner et al., 2014).

Their early divergence in the insect phylogenetic tree, the occurrence of different reproductive systems and ploidy levels and their patterns of distribution make bristletails in the genus *Machilis* an interesting study object in ecology and evolution. Here, we provide novel genomic resources for this system and we discuss the evolutionary history of an Alpine bristletail species. We present the first de-novo assembled transcriptome obtained from 17 triploid individuals of *Machilis pallida* from six populations in the European Alps as well as a de-novo assembled mitochondrial genome. We then utilize these new resources to combine mitochondrial variation as well as common and rare nuclear variation and assess the distribution of genetic diversity among those sets of markers to elucidate the evolutionary history of this species. Lastly, we discuss our findings in the context of current hypotheses on the reproductive mode, ploidy and distribution of *M. pallida*.

## 2. Material and Methods

### 2.1. Sample collection

We collected 17 adult female (the only sex found so far) *Machilis pallida* specimens at six locations in the eastern Alps (Laempermahdspitze (L, n = 4 individuals), Kesselspitze (K, n = 1), Padasterjochhaus (P, n = 5), Obernberger Tribulaun (O, n = 2), Murmeltierhuette (M, n = 2) and Grosté Seilbahn Bergstation (G, n = 3)) in September 2016 (Fig. 1, Tables 1 and S1). This sampling design includes putative central nunatak populations (L, K, P, O) and putative peripheral nunatak populations (M and G). All individuals were transported alive to the Department of Ecology at the University of Innsbruck, Austria and kept in plastic boxes supplemented with humid gravel from the collection site at 10 °C and a 12:12 hours light:dark cycle for four days to normalize gene expression.

**Figure 1.**
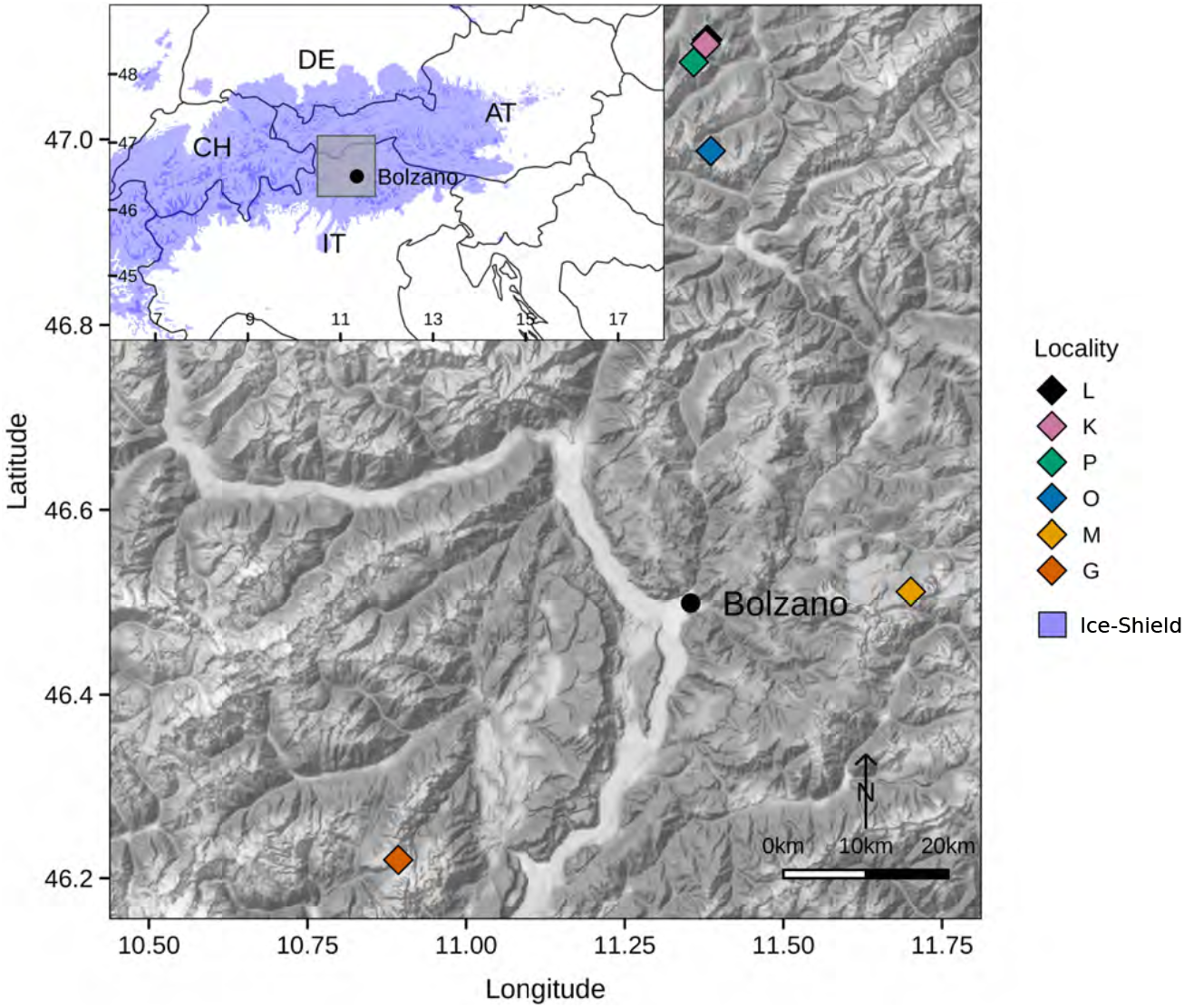
Sampling locations of the six populations of *Machilis pallida*. Colors of diamonds correspond to the different populations; Laempermahdspitze (L, black), Kesselspitze (K, pink), Padasterjochhaus (P, green), Obernberger Tribulaun (O, blue), Murmeltierhuette (M, yellow) and Grosté Seilbahn Bergstation (G, orange). The overview map shows the extent of the Alpine glaciers during the last glacial maximum.

**Table 1.** Sample information. Locality abbreviations correspond to Laempermahdspitze (L), Kesselspitze (K), Padasterjochhaus (P), Obernberger Tribulaun (O), Murmeltierhuette (M) and Grosté Seilbahn Bergstation (G).

| ID | Individual | Locality | Longitude | Latitude | Country | Elevation (m a.s.l.) |
| --- | --- | --- | --- | --- | --- | --- |
| 1 | 92360 | K | 11.377325 | 47.102333 | AT | 2280 |
| 2 | 92361 | L | 11.378703 | 47.107025 | AT | 2220 |
| 3 | 92362 | L | 11.379536 | 47.108000 | AT | 2220 |
| 4 | 92363 | L | 11.379769 | 47.106950 | AT | 2220 |
| 5 | 92364 | L | 11.380089 | 47.107153 | AT | 2220 |
| 6 | 92365 | O | 11.385581 | 46.987497 | AT | 2000 |
| 7 | 92366 | O | 11.385383 | 46.987583 | AT | 2000 |
| 8 | 92367 | G | 10.893125 | 46.219956 | IT | 2400 |
| 9 | 92368 | G | 10.889464 | 46.222336 | IT | 2400 |
| 10 | 92369 | G | 10.892083 | 46.220806 | IT | 2400 |
| 11 | 92370 | M | 11.700775 | 46.511847 | IT | 2200 |
| 12 | 92371 | M | 11.699558 | 46.511008 | IT | 2200 |
| 13 | 92372 | P | 11.358669 | 47.082906 | AT | 2320 |
| 14 | 92373 | P | 11.358606 | 47.083008 | AT | 2320 |
| 15 | 92374 | P | 11.358481 | 47.082889 | AT | 2320 |
| 16 | 92375 | P | 11.358458 | 47.082933 | AT | 2320 |
| 17 | 92376 | P | 11.358381 | 47.082917 | AT | 2320 |

### 2.2. RNA extraction, library preparation, and sequencing

Individuals were shock-frozen in liquid nitrogen and we performed RNA extraction using the Macherey and Nagel (Düren, Germany) Nucleospin RNA kit following the instructions of the manufacturer. Library construction and sequencing were performed by IGATech (Udine, Italy). We then constructed a barcoded library for all individuals with the Illumina (San Diego, United States of America) TruSeq Stranded mRNA Library prep kit following the manufacturer’s instructions. Libraries were then sequenced on an Illumina HiSeq 2500 platform in 250 bp paired-end rapid run mode.

### 2.3. Transcriptome assembly and mitochondrial reads

We performed quality control of the raw data with FastQC (Andrews, 2010)before concatenating raw reads from the 17 individuals. BBtools v37.36 (Bushnell, 2021) was used for (i) quality trimming and quality filtering (bbduk.sh with minimum length of 10), (ii) decontamination (with bbduk.sh and reference phix174_ill.ref.fa, k-mer size of 31 and hamming distance of 1), and (iii) read normalization (with bbnorm.sh and read coverage of 60 and minimum depth of 5). Quality-trimmed and normalized reads were assembled using Trinity v2.2 (Grabherr et al., 2011) with the following settings: strand-specific RNA-Seq read orientation, minimum contig length of 500, Jaccard clip, and normalization of reads. After assembly, we performed cleaning steps to remove contaminants and redundancy in the transcriptome: (i) Blobtools v1.0 (Kumar et al., 2013) was used for contaminant filtering with two Blastn (Altschul et al., 1990) e-value cutoffs of 1e-3 and 1e-5 to identify contaminants. In this step, transcripts of viruses, archaea, bacteria, and fungi were removed from the dataset. (ii) CD-HIT-EST v2 (Fu et al., 2012) was used to find unigenes (options: alignment coverage of 0.9, word length of 8, and cluster to most similar cluster (g) of 1). When multiple transcripts had the same BLAST hit, only the longest was retained. After each cleaning step, a completeness check was performed using Universal Single-Copy Orthologs (BUSCO) software v3 (Simão et al., 2015). Further, we calculated Nx, ExN50 statistics, and the percentage of raw reads using Trinity v2.2 and we estimated read abundance with the package RSEM (Li and Dewey, 2011). Finally, we aligned individual raw reads to the assembled transcriptome using Bowtie2 (Langmead and Salzberg, 2012). With Transdecoder v2.0.1 (Haas et al., 2013), possible open reading frames (ORFs) were detected. With these predictions, the annotation with Trinotate v3.1.1 (Bryant et al., 2017) was performed by doing a homology search in BLAST (Altschul et al., 1990) and Swissprot (The UniProt Consortium, 2017), protein domain identification with HMMER v3 (Finn et al., 2011) and PFAM v31.0 (Finn et al., 2014), protein signal peptide prediction with signalP v4.1 (Petersen et al., 2011), transmembrane domain prediction with tmHMM v2.0c (Krogh et al., 2001) and further annotation databases (eggnogg; (Huerta-Cepas et al., 2016), Gene Ontology (GO; (Ashburner et al., 2000), and Kegg (Kanehisa et al., 2012)). GO terms per gene were visualized with Web Gene Ontology Annotation Plot v2.0 (WEGO; (Ye et al., 2006)).

We further created a de-novo mitochondrial assembly of *M. pallida* using MITObim (Hahn et al., 2013), which employs a mitochondrial baiting and iterative mapping approach using the MIRA assembler (Chevreux et al., 1999). We used a kmer length of 31 for bait fishing with mirabait using genome skimming data (individual 92010, Murmeltierhuette, ERS4357532; SAMEA6593248; T. Dejaco, unpubl.) and the *Songmachilis xinxiangensis* mitochondrium as baiting sequence (He et al., 2013) (with MIRA v4.0.2 (Chevreux et al., 1999)) with 30 iterations. We used ORFfinder to search for insect mitochondrial ORFs in the mitochondrial contig, and from the resulting ORFs, we blasted the 10 longest hits using smartBLAST (NCBI, 2021).

### 2.4. Alignment and variant calling

We used bbmap (Bushnell, 2021) to align the trimmed and decontaminated reads of all 17 individuals with a minimum identity score of 0.97 for the alignments to the de-novo assembled transcriptome, and a minimum identity score of 0.90 for the alignments to the mitochondrium. After removing unmapped reads for both nuclear and mitochondrial alignments with samtools v1.9 (Li et al., 2009), we marked duplicate reads with Picard v2.19.1 (Broad Institute, 2018). For the nuclear RNA, we called variants with GATK v3.8 and the UnifiedGenotyper tool (McKenna et al., 2010). Specifically, for nuclear RNA reads, we called variants as triploid, with a minimum phred-scaled confidence threshold for variants to be called of 50, three alternative alleles and the SNP genotype likelihood model. For the mitochondrial reads, we used bbmap to call variants with quality score recalibration (bbvarMT.sh) and a ploidy of two. After calling variants, we filtered both the nuclear and mitochondrial variants with a minimum coverage of 128, a minimum mapping quality of 50 and minimum occurrence of four sequences with the alternative allele. Additionally, we only kept substitutions that were not fixed for either the reference or alternative allele. After filtering the variants, we split the nuclear variants into common and rare variants, where variants with allele frequency < 0.1 were considered rare.

### 2.5. Population genetic and phylogenetic analyses

We extracted genotypes for mitochondrial variants, as well as common and rare variants using custom python scripts (mafFltr.py). We then calculated Hamming distances from the diploid (mt) and triploid (common and rare) variants for Multidimensional Scaling (hereafter referred to as Principal Coordinates Analysis, PCoA) in R (R Core Team, 2020). We further converted the genotypes of all three variant types into Nexus format (vcf2nex.py) to compute distance matrices and obtain neighbor-joining (NJ) trees for all three variant types in R using the ape package (Paradis and Schliep, 2019), and to obtain haplotype networks for rare and mitochondrial variants using the pegas package (Paradis, 2010).

## 3. Results

### 3.1. Transcriptome assembly and mitochondrial reads

The mean read number per individual after quality trimming, decontamination and normalization amounted to 8 million (M) reads (sd = 1.7 M) (see Table S1), resulting in a total of 359 M reads used for the assembly. After quality filtering, decontamination, and normalization, 46,748,840 reads were used for the assembly. In total, 289,342 contigs (Trinity transcripts) and 117,970 genes (Trinity genes) were assembled. After removal of redundancy in the dataset, the final set contained 159,192 contigs and 106,399 genes. GC content was 40.39% (Table 1), N50 contig length 1,959 bp, and E80N50 contig length 2,535 bp. The overall read alignment was 80.07%, of which 68.95% were properly paired. BUSCO revealed a high completeness of the transcriptome (93.3%) when probing the insect database.

In the 159,192 transcripts, 124,989 ORFs were detected. Overall, 61,186 transcripts (64% of all assembled transcripts) were annotated. Of these, 50% were assigned to a unique protein in UniprotKB using Blastx. With Blastp, 38% of the transcripts showed a protein hit. In Eggnog, Kegg, and Blast GO terms, 34%, 35%, and 43% of the transcripts showed a hit, respectively. A total of 65,782 GO terms were assigned to the transcripts. Of these, approximately 32% were described by the aspect biological process, 34% by cellular component, and 34% by their molecular function (Table S2, https://github.com/mphaider/M.pallida.git).

In the *M. pallida* mitochondrial genome assembly, MITObim reached a stationary read number of 54,161 reads after 25 iterations of baiting and mapping, with a length of 15,836 bp for the resulting mitochondrial contig. The ten longest ORFs on the mitochondrium are summarized in Table S3.

### 3.2. Alignment and variant calling

Across the 17 *M. pallida* individuals, the mean number of reads amounted to 40.97 M (sd = 8.7 M) after quality trimming and decontamination. Reads mapped with an average rate of 69.67% (sd = 1.46%) and 7.01% (sd = 2.15%) to the nuclear and the mitochondrial assembly, respectively. After variant calling and filtering, we found a total of 213,321 variants (from 1,196,769 unfiltered variants) using the nuclear assembly as reference, of which 201,195 variants were common and 12,126 variants were rare (i.e. 5.69% of variants were lower than an allele frequency of 0.1). For the mitochondrial alignments, we found 29 variants after filtering (with 32 unfiltered variants).

### 3.3. Population genetic and phylogenetic analyses

Mitochondrial variants were relatively well-resolved in the PCoA, where the three highest Principal Coordinates (PCos) explained 94,5% of the overall variance across the 29 variants (Fig. 2 and S1). We found a tight central cluster formed by individuals from O, L, and K, which was adjacent to all individuals from the P site. As shown in Figure 2, M and G locations are placed farthest from the central cluster, and they are positioned on opposite sides of the central cluster (i.e. the two southern locations (M and G) are both closer to the central cluster than to each other, differentiated on all three PCos).

**Figure 2.**
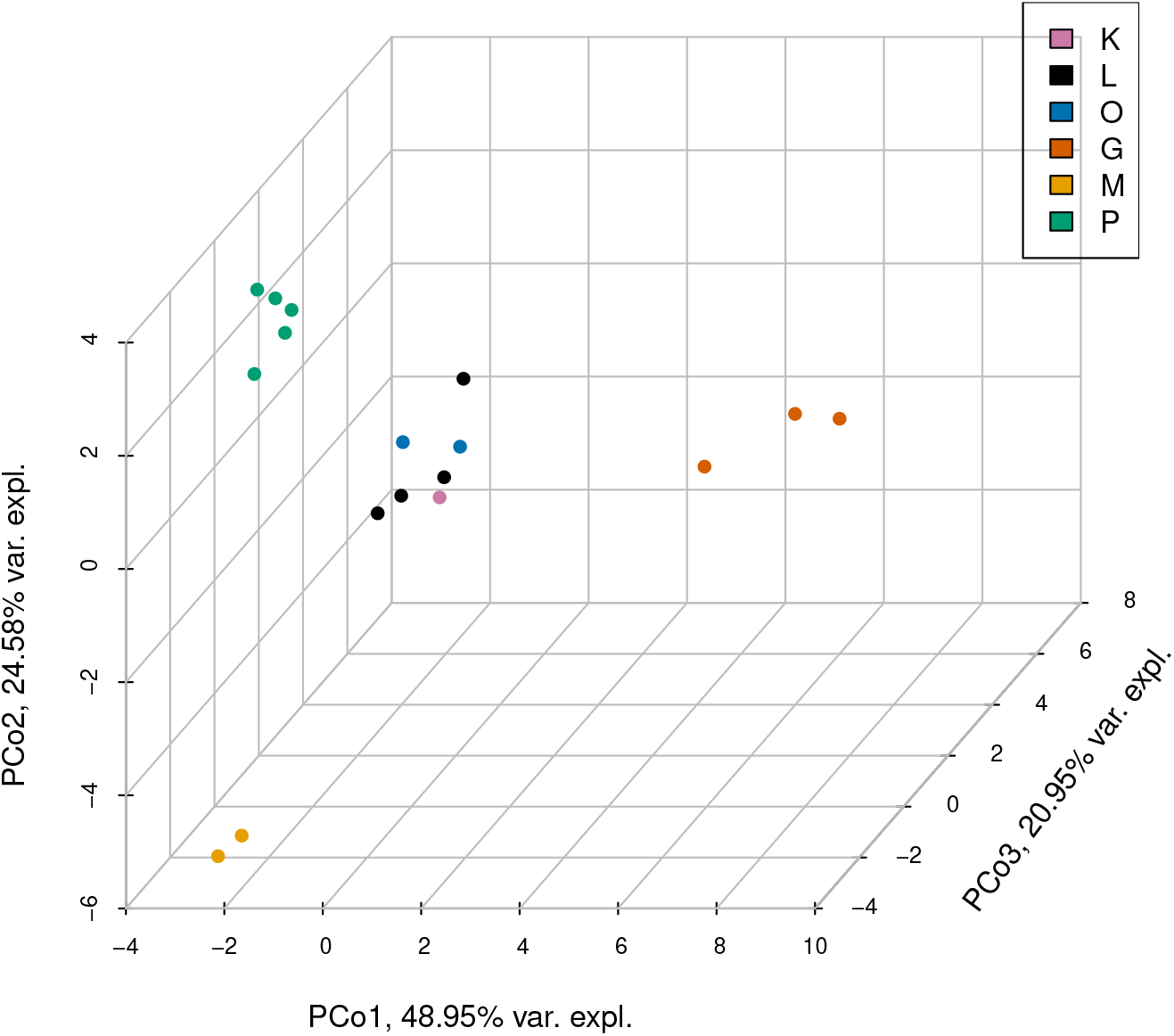
Principal coordinate analysis of 29 mitochondrial variants. Colors correspond to the different populations; Laempermahdspitze (L, black), Kesselspitze (K, pink), Padasterjochhaus (P, green), Obernberger Tribulaun (O, blue), Murmeltierhuette (M, yellow) and Grosté Seilbahn Bergstation (G, orange).

The first three PCos for 190,858 common nuclear variants explained about 27% of the overall variance found and are shown in Figure 3 (see also Fig. S2). Here, we also see a tight central cluster, formed by the sites of L, O, two samples from G as well as one individual from M. Close by, P again forms a distinct cluster, yet close to the central group (mainly distinguished on PCo 1 and PCo 3). The remaining M individual is far removed (distinguished mainly on PCo 1), and the last two individuals (from K and G) are somewhat close to each other, yet far removed from the rest, distinguished mainly on PCos 1 and 3. Note that PCos 1 and 3 share patterns, with PCo 2 seemingly differing based on the variance encountered.

**Figure 3.**
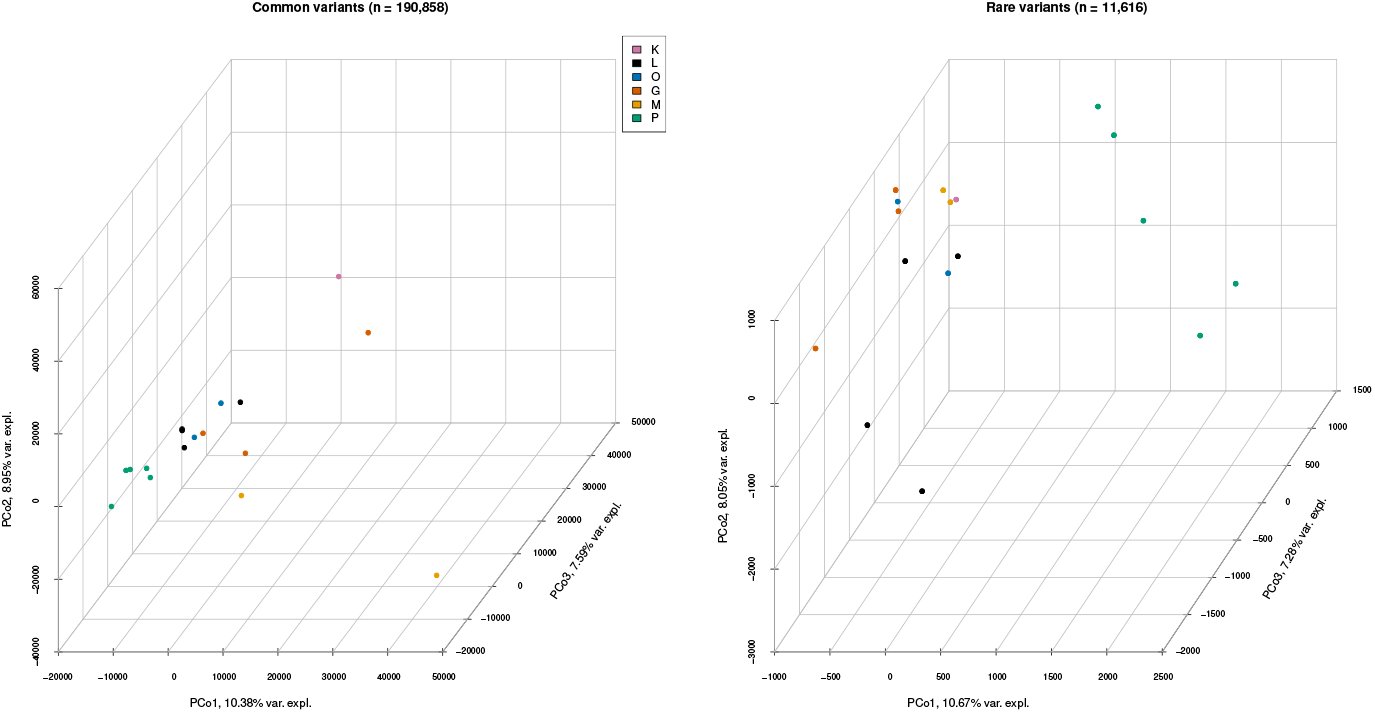
Principal coordinate analysis of common (left panel) and rare (right panel) variants with 190,858 and 11,616 variants. Colors correspond to the different populations; Laempermahdspitze (L, black), Kesselspitze (K, pink), Padasterjochhaus (P, green), Obernberger Tribulaun (O, blue), Murmeltierhuette (M, yellow) and Grosté Seilbahn Bergstation (G, orange).

Principal coordinates of 11,616 rare nuclear variants explained about 26% of the overall variance across the 17 individuals. Figures 3 and S3 depict a cluster, formed by K, M, two samples from G, and two individuals from L. The remaining individual from G is close to the remaining individuals from G on PCo 1 and 2 but different on PCo 3. The remaining two L individuals were predominantly distinguished on PCos 2 and 3. Further, variants of P spread the five individuals mainly across PCos 1 and 3.

In the mitochondrial haplotype network, we found 11 haplotypes (Fig. 4), where the two Southern localities (G and M with haplotypes g, h, and i) both split off from a central batch, containing one haplotype each from L and O (e), where link lengths to the Southern haplotypes consist of at least five steps. All remaining haplotypes fork off from the central haplotype (e), with three more haplotypes for L (b, c, and d), one for K (a), two for P (j and k), and one more haplotype for O (f), which links through a haplotype from L (b). A haplotype network for rare variants showing 17 haplotypes for 17 individuals can be found in the Supplementary Material (Fig. S4).

**Figure 4.**
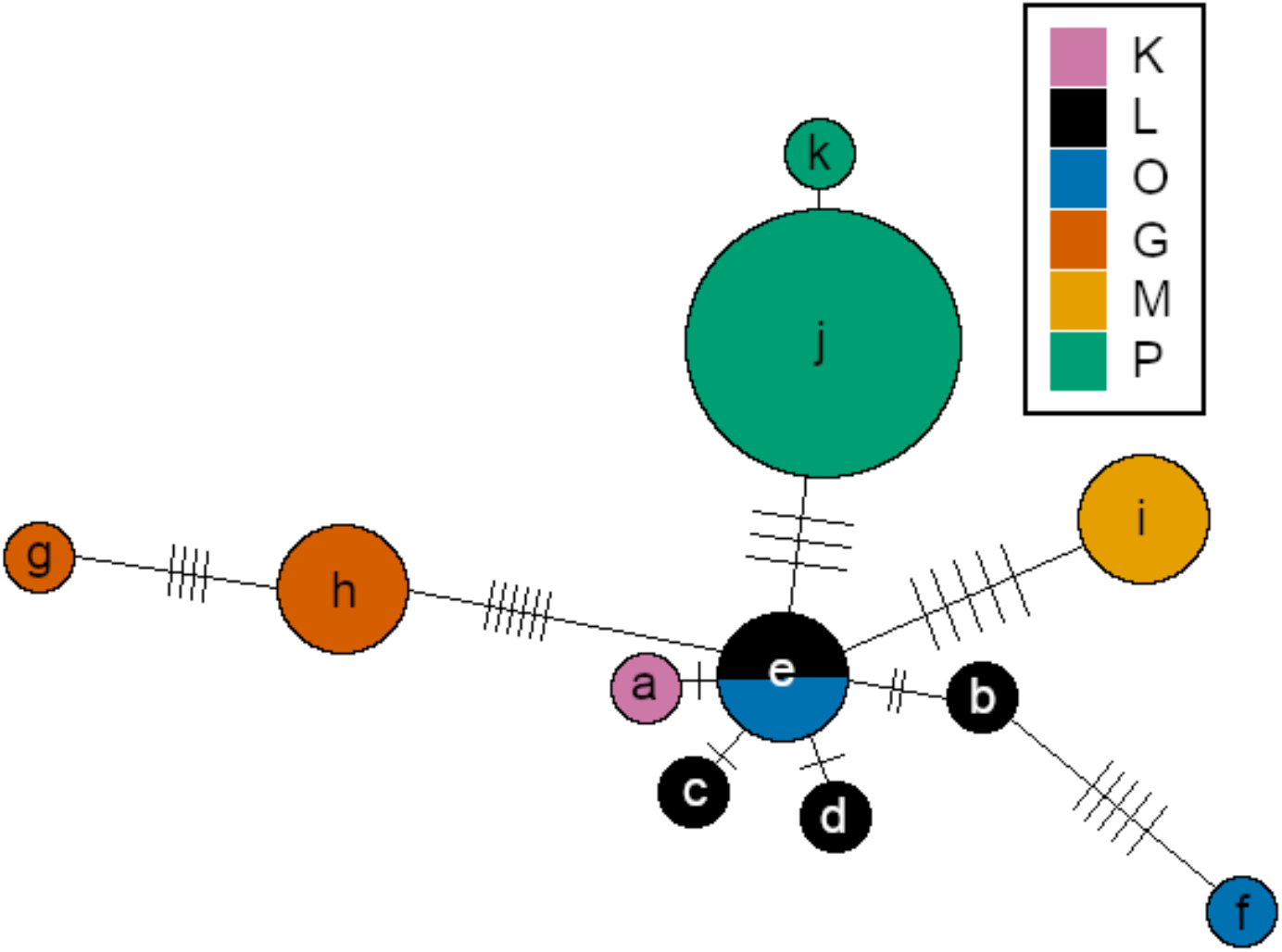
Haplotype network for mitochondrial variants (n = 29). Note that a link length of two between haplotypes a and d was omitted in this figure. Colors correspond to the different populations; Laempermahdspitze (L, black), Kesselspitze (K, pink), Padasterjochhaus (P, green), Obernberger Tribulaun (O, blue), Murmeltierhuette (M, yellow) and Grosté Seilbahn Bergstation (G, orange). Lowercase letters a to k denote the 11 mitochondrial haplotypes.

The resolution of the neighbor-joining trees varied substantially among the three variant types. For the 29 mitochondrial variants, the Northern locality of P falls into one polytomous group, and so do the Southern localities of M and G as shown in Figure 5. The remaining individuals, from localities L, K, and O, are not resolved. The 12,126 rare nuclear variants do not resolve the relationships among the individuals and we observe a single polytomous block. The highest level of resolution, however, was achieved with the 201,105 common variants, revealing a clear distinction into localities G and M, the central group of L, O, and K, and finally the polytomous individuals from locality P.

**Figure 5.**
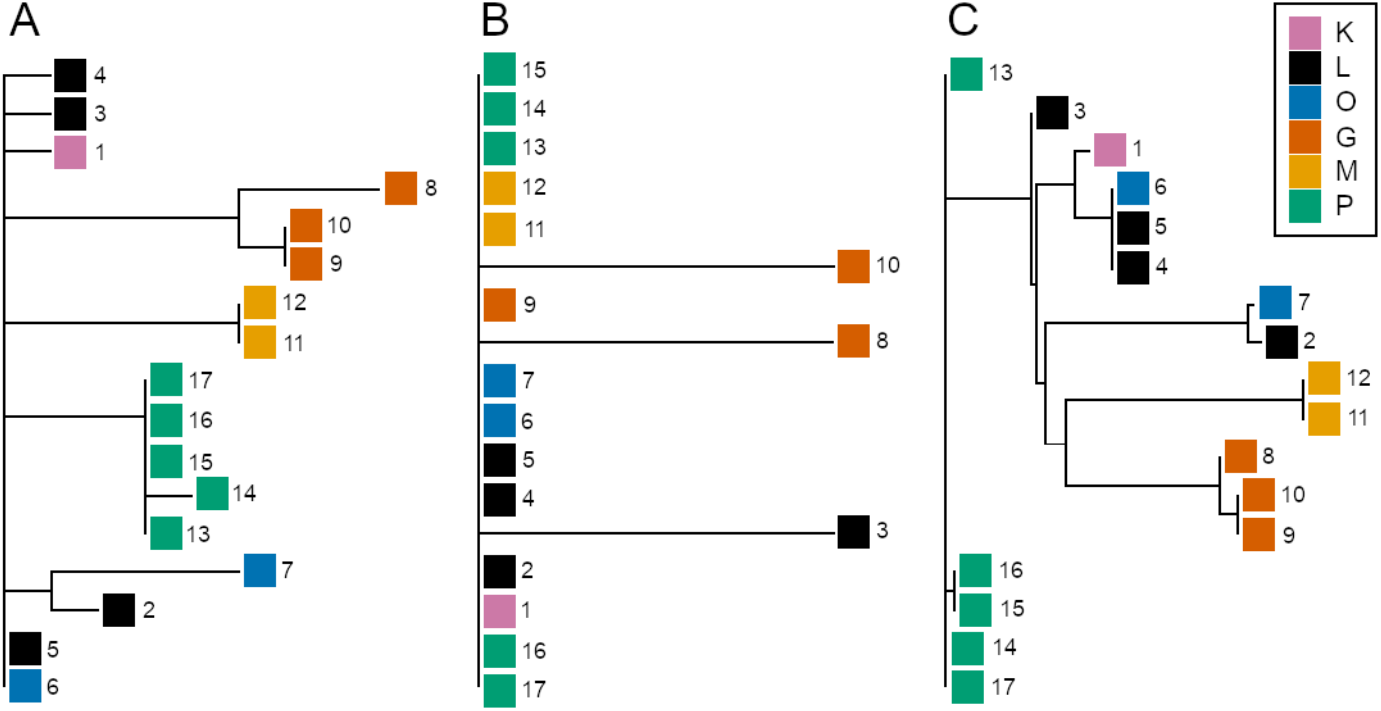
Neighbour-joining trees for all three variant types, with A. Mitochondrial variants (n = 29), B. Rare variants (n = 12,126), and C. Common variants (n = 201,105). Colors correspond to the different populations; Laempermahdspitze (L, black), Kesselspitze (K, pink), Padasterjochhaus (P, green), Obernberger Tribulaun (O, blue), Murmeltierhuette (M, yellow) and Grosté Seilbahn Bergstation (G, orange).

## 4. Discussion

This work contributes to ongoing efforts directed towards a better understanding of the complex evolutionary history of Alpine bristletails. For this purpose, we established new resources, particularly novel transcriptome and mitochondrial assemblies for *Machilis pallida* Janetschek, 1949. Together with various genotyping-by-sequencing approaches, transcriptomic and mitochondrial sequence data represent cost-effective options for assessing patterns of genetic diversity (Hahn et al., 2013; Hirsch et al., 2014), in particular in non-model organisms where a reference genome is absent and/or genomes are very large (i.e. >1 Gb in *Machilis* (Gassner et al., 2014)). In addition, RNA-seq data and transcriptome assemblies are valuable resources for the functional annotation of a reference genome, which can be produced at a later stage.

Using nuclear and mitochondrial RNA-seq data for 17 individuals from six sampling localities, we assess different types of variation – mitochondrial as well as rare and common nuclear variants – to capture and reconstruct different aspects of the evolutionary history of *M. pallida* and reassess previous work. In particular, the suggested parthenogenetic mode of reproduction (Wachter et al., 2012) and polyploidy (Gassner et al., 2014) in *M. pallida* certainly affect our expectations regarding the patterns of genetic diversity for the different kinds of variation under various evolutionary scenarios.

Population genetic theory predicts that asexual populations should 1) have lower effective population sizes (Barton and Charlesworth, 1998), and 2) be less efficient in their adaptive potential, since beneficial mutations would be lost more easily in asexual compared with sexual populations (Muller, 1964). A beneficial mutation needs to confer a strong selective advantage to avoid loss through clonal interference and Muller’s Ratchet (Fisher, 1930; Muller, 1932, 1964; Hill and Robertson, 1966; Felsenstein, 1974; Charlesworth and Charlesworth, 1997).

However, transiently abundant beneficial mutations that do not go to fixation might be common in asexual populations, which might experience a leapfrog effect, where the common genotype is less closely related to the immediately preceding common genotype, but more closely related to earlier genotypes (Gerrish and Lenski, 1998). In other words, two individuals sampled from different asexual populations might appear to be more closely related to each other than to individuals from their own population if there is more than one asexual lineage present therein.

Further, it is predicted that in asexual diploid species, alleles on chromosome homologues would show independent evolution (referred to as the ‘Meselson effect’ sensu Birky 1996, Welch and Meselson 2000, Butlin 2002), but it is not clear whether this is necessarily the case in organisms with higher ploidy levels.

Polyploidy is frequently coupled with asexuality (Otto and Whitton, 2000) and this combination presents particular challenges for the interpretation of genomic data. In general, polyploid asexual species are assumed to be relatively rare (White, 1973; Burch and Jung, 1993; Dufresne and Hebert, 1994; Otto and Whitton, 2000), and most obligately asexual lineages in plants and animals have evolved relatively recently (Maynard Smith, 1978). Moreover, it was found that polyploid populations more often tend to have multiple rather than single origins in plants (Soltis et al., 1993; Soltis and Soltis, 1999) and in animals (reviewed in Otto and Whitton 2000, see also Chaplin and Hebert 1997, Dufresne and Hebert 1994). Multiple origins of polyploidy are thought to arise via either a high rate of polyploidization during initial establishment of a given lineage and/or recurrent gene flow with related diploid taxa (Otto and Whitton, 2000).

Based on the assumptions stemming from aforementioned theoretical work, we will now discuss the significance of the patterns found in the different variant types examined in this study. Notably, the analyses of the three different variant types revealed distinct patterns suggesting that they indeed capture different aspects of the evolutionary history of *M. pallida*. Consistent with findings by Wachter et al. (2012), the mitochondrial data mostly reflect geography, with the two southern localities and populations from the Alpine main ridge representing three main groups (Fig. 2 and S1). Within the central Alpine populations, however, P forms a coherent, somewhat delimited cluster. While localities K and L are very close to each other and connected by a ridge, P is demarcated by a valley and O is farthest away and separated by a larger valley. Therefore, the P cluster may, at least partly, also mirror a geographic pattern. However, P is the locality with the highest number of samples, and we cannot exclude a sample size bias in our analyses.

The common nuclear variants show a more heterogeneous picture but are broadly consistent with the mitochondrial results confirming the differentiation of the geographically separated central Alpine and southern localities as well as the P cluster. We note, however, that the differentiation between North and South is less clear and more gradual (Fig. 3 and S2). Moreover, we do not find support for the existence of two main nuclear clusters suggested by an earlier admixture analysis on AFLP data (Wachter et al., 2012) (discussed below).

The results from the rare nuclear variants are in stark contrast to the other two marker types. We do not find any obvious patterns mirroring populations or geographic distances with the sole exception of the P cluster in the PCoA, which may be driven by sample size (Fig. 3 and S3).

For the interpretation of the observed diversity patterns, we consider four simplistic scenarios that differ in whether parthenogenesis emerged before or after populations split and in the presence and absence of migration (Fig. 6). Assuming a single origin of parthenogenesis in an unstructured population followed by immediate dispersal of this lineage to the current locations (Fig. 6 A and C), we would not expect to see clustering by geography. Later migration between nearby populations (Fig. 6 D) could, however, create a signature of geography and isolation by distance and, for example, explain the coherence of the central Alpine cluster. *Machilis* species are considered slow dispersers based on the fact that they lack wings and due to the high degree of endemism in the genus. However, their actual dispersal potential has not been quantified so far (Sturm and Machida, 2001, p. 61/62). A related scenario would involve a single origin of parthenogenesis followed by an extended time period allowing for range expansion and differentiation of the parthenogenetic lineage and subsequent fragmentation, yielding the current distribution pattern.

**Figure 6.**
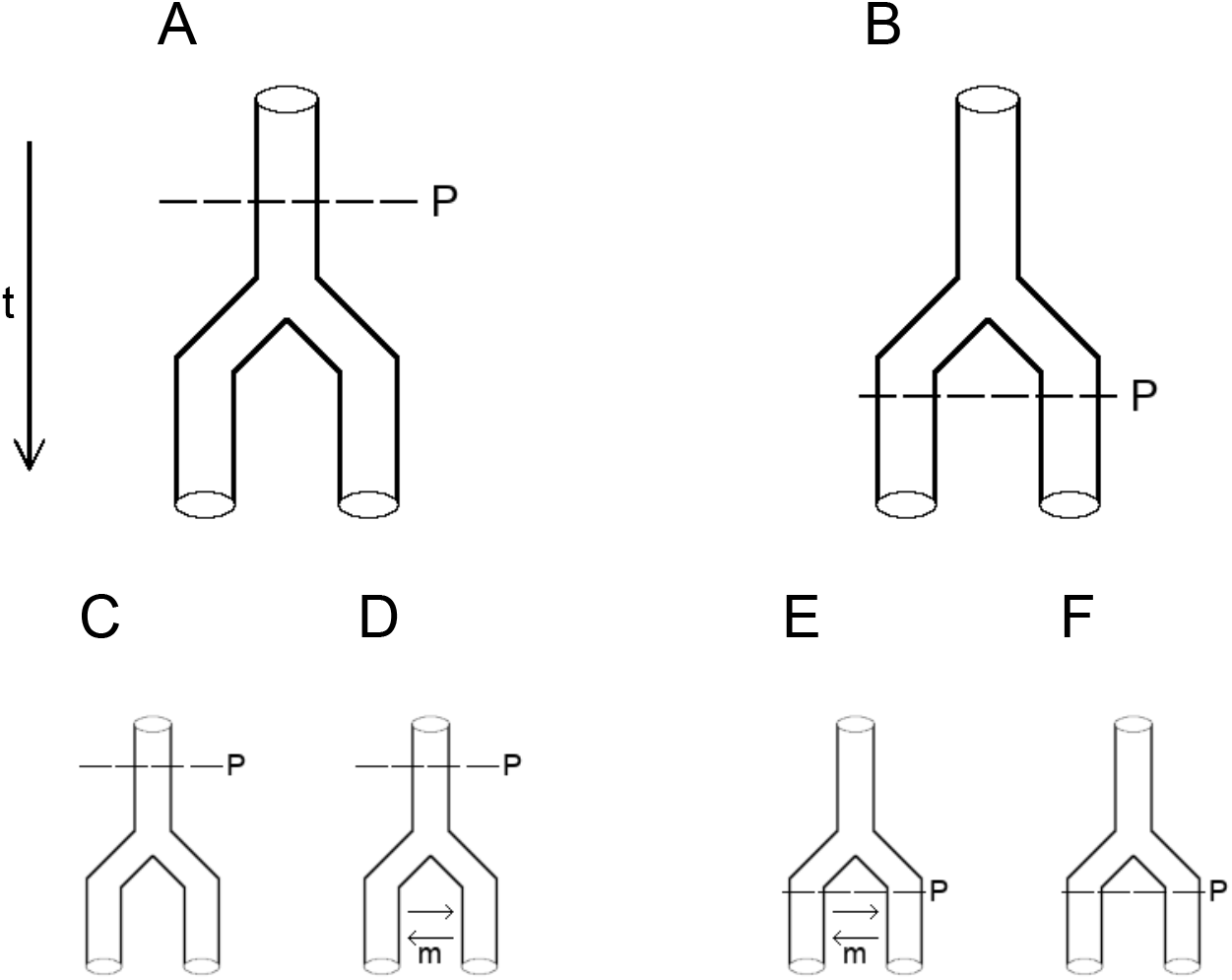
Hierarchical representation of simplified putative scenarios of the evolutionary history of *Machilis pallida* that differ in the timing of the onset of parthenogenesis (A, B) and the presence of migration (C-F).

Alternatively, the clusters visible in the mitochondrial and common nuclear data may reflect population structure already present in a more widespread sexual progenitor, whereas the lack of structure in the rare nuclear variants may be attributable to the presence of asexual lineages as well as the small sample sizes for single populations. In this scenario, multiple transitions to parthenogenesis after the build-up of population structure need to be invoked (Fig. 6 B, E, and F) but it would not be necessary to postulate migration to explain the geographic clusters (Fig. 6 E and F). The fact that the mitochondrial variants show more distinct clusters compared with the common nuclear variants is consistent with the smaller effective population size of mitochondrial DNA and therefore a stronger effect of drift. Alternatively, common nuclear variants could exhibit the aforementioned leapfrog effect (Gerrish and Lenski, 1998). That is, if enough common variants show a pattern, where an individual appears more closely related to a conspecific from a different population, then this could lead to common nuclear variants exhibiting a different picture than rare nuclear or mitochondrial variants, where individuals from different populations form tight clusters. However, while common nuclear variants do in fact show such a pattern, we cannot make a definitive statement as to the underlying processes. While we focus on a few simple scenarios here, we note that arbitrarily complex scenarios including combinations of multiple, temporally separated origins of parthenogenesis, rare sexual reproduction, colonization and extinction cycles, and stepping stone models could also fit our data.

The observed patterns of genetic diversity might be consistent with ice age survival of *M. pallida* on central and/or peripheral nunataks. The central position of the populations located at the main ridge of the Alps relative to the two southern populations in the mitochondrial haplotype network (Fig. 4) could indicate dispersal from a central Alpine refugium. However, as mentioned before, there could be different scenarios, congruent with the observed data. For instance, the neighbor-joining tree constructed from common nuclear variants (Fig. 5 C) proposes a closer relationship between the two southern populations consistent with the existence of central and peripheral refugia. Additionally, the aforementioned leapfrog effect may also affect the observed relationships between populations and thus, interpreting these patterns warrants caution.

Wachter et al. (2012) proposed central and peripheral nunatak survival based on the geographic distribution of mitochondrial COI haplotypes and two nuclear AFLP clusters across three sampling localities. Under the assumption of slow and limited dispersal of this apterygote insect species, our data could be explained by such a scenario. In contrast to these previous results, however, we find no evidence for the presence of two distinct nuclear clusters but instead reveal a more complex population structure. This may be due to differences in marker types and sample size or due to the k=2 conundrum (Janes et al., 2017), the tendency of the deltaK method (Evanno et al., 2005) to frequently identify k=2 as top hierarchical level in STRUCTURE analyses (Pritchard et al., 2000). Moreover, our strategy of leveraging the information contained in sets of different variants highlights possible, more complex variations of this simplistic scenario as outlined above despite small sample size. Therefore, applying this approach to a larger sampling scheme, both in terms of populations and individuals, combined with simulation studies holds great potential to fully elucidate the distribution and migration patterns during the ice ages, rigorously test competing hypotheses and date dispersal events as well as the onset of parthenogenesis.

Polyploidization is highly associated with parthenogenetic reproduction in animals and also frequently correlated with hybridization (Otto and Whitton, 2000). It is conceivable that polyploidization events were accompanied by transitions to parthenogenesis in *M. pallida* and that hybridization was involved (e.g. Dejaco et al., 2016). Moreover, since multiple origins of polyploidy are not uncommon (reviewed in Otto and Whitton, 2000; Soltis et al., 1993; Soltis and Soltis, 1999; Chaplin and Hebert, 1997; Dufresne and Hebert, 1994), the putative joint occurrence of parthenogenesis and polyploidy would neither contradict scenarios involving a single or multiple origins of parthenogenesis. To actually assess these relationships, however, additional work is required.

## 5. Conclusions

In this study, we assess genetic diversity patterns for different variant types in a parthenogenetic, triploid, endemic Alpine bristletail species. We demonstrate that mitochondrial and common nuclear variants mirror geographic patterns. Moreover, we highlight that different types of variants capture different aspects of the evolutionary history of the species and outline the potential of their combined consideration for unraveling more complex scenarios. We emphasize that *M. pallida* and the genus *Machilis* in general represent an interesting study system for the evolution of different sexual strategies, polyploidization and genome re-organisation, as well as for adaptation to environmental change. We also highlight the need for further studies and the presented novel resources, transcriptome and mitochondrial assemblies together with transcriptome data for 17 individuals, will facilitate future research in this diverse system.

## Abbreviations

mtDNA: mitochondrial DNA
ptDNA: plastid DNA
PCoA: Principal Coordinates Analysis

## 6. Data Availability

Data generated in this study are deposited at the European Nucleotide Archive (ENA) under the study accession ERP120116. Details on sample and raw sequence data accessions can be found in Table S4 and transcriptome and mitochondrial assembly ENA accessions are ERZ1673957 and ERZ1668275. Details on the de-novo transcriptome assembly workflow and variant calling as well as Table S2 are deposited at https://github.com/mphaider/M.pallida.git. Code for mitochondrial variant calling and population genetics analyses are deposited at https://github.com/schimar/mpallida19.git.

## 7. Acknowldegements

We thank Melitta Gassner and Richard Hastik for collecting the individuals used in this study, Philipp Andesner for support in the wetlab, Thomas Dejaco for sharing his expertise on the study system and genome skimming data, and Federica Cattonaro for the informed execution of the sequencing order. The computational results presented here have been achieved in part using the MACH2 Interuniversity Shared Memory Supercomputer, the HPC infrastructure of the University of Innsbruck and Vienna Scientific Cluster (VSC). We thank Hermann Schwärzler and Francesco Cicconardi for bioinformatics support. This research was funded in part by the Austrian Science Fund (FWF):P30861 and the Autonomous Province of South Tyrol [project-ID: 1/40.3; 27 January 2014].

