## Supplementary Material for "Evolutionary history of an Alpine archaeognath (*Machilis pallida*) – insights from different variant types"

---

---

*Corresponding author

*Email addresses:* (Marlene Haider), (Martin P. Schilling), (Markus H. Moest), (Florian M. Steiner), (Birgit C. Schlick-Steiner), (Wolfgang Arthofer)

[1]equally contributing first author

[2]equally contributing senior author

### 1. Supplementary Figures

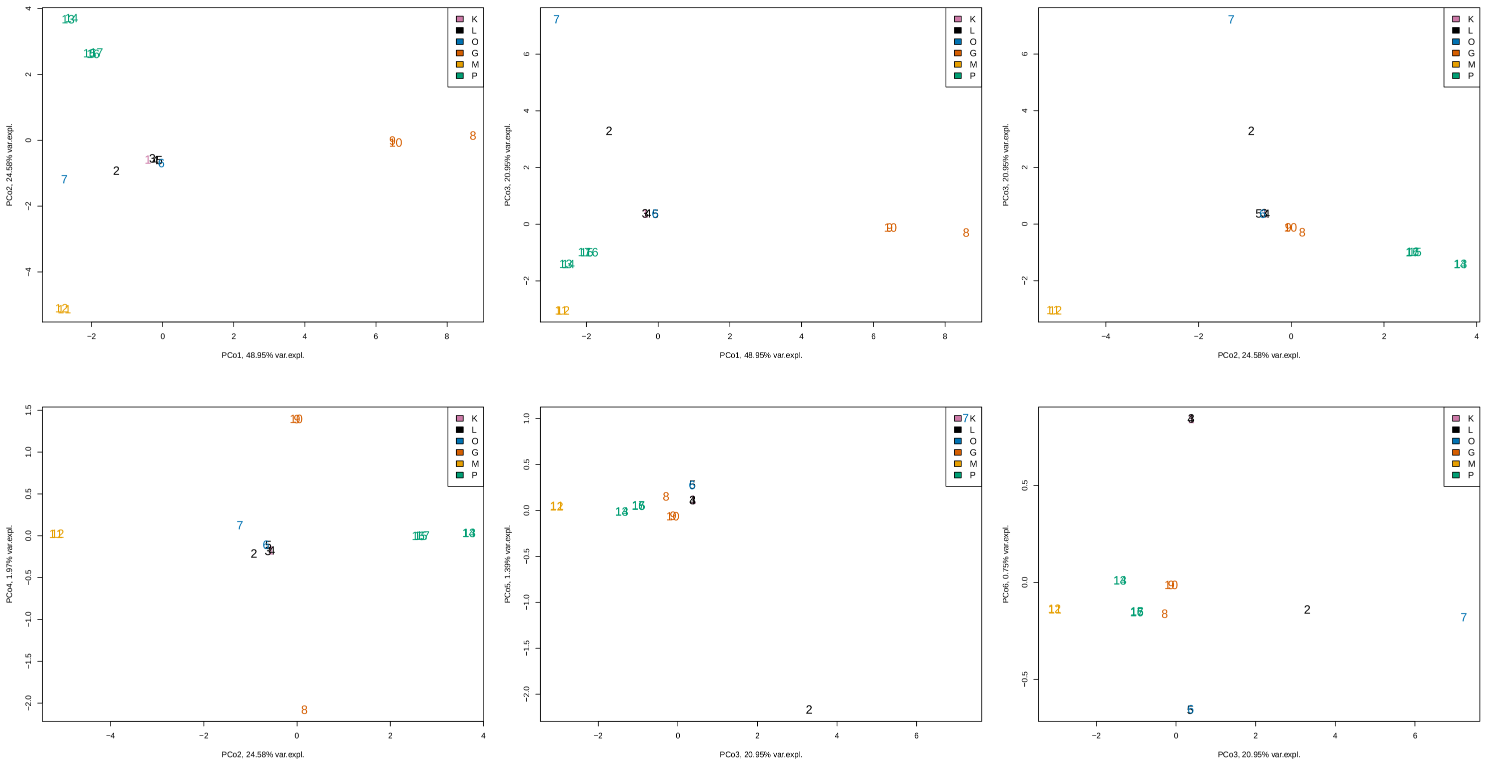


**Figure S1.** Principal coordinate analysis of 29 mitochondrial variants. Combinations of the first six principal coordinate axes are shown. Colors correspond to the different populations; Kesselspitze (K, pink), Laempermahdspitze (L, black), Obernberger Tribulaun (O, blue), Grosté Seilbahn Bergstation (G, orange), Murmeltierhuette (M, yellow), and Padasterjochhaus (P, green).

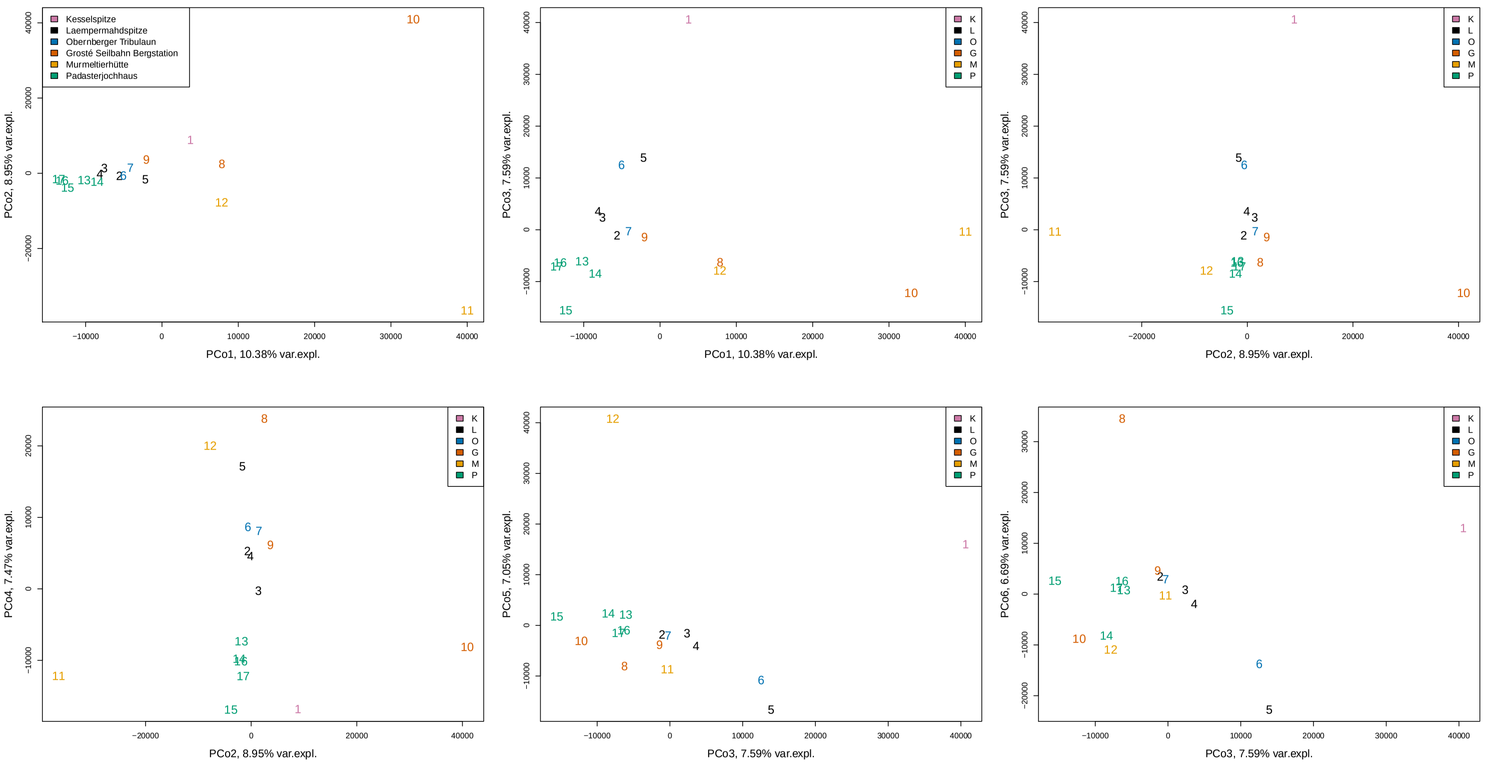


**Figure S2.** Principal coordinate analysis of 190,858 common nuclear variants. Combinations of the first six principal coordinate axes are shown. Colors correspond to the different populations; Kesselspitze (K, pink), Laempermahdspitze (L, black), Obernberger Tribulaun (O, blue), Grosté Seilbahn Bergstation (G, orange), Murmeltierhuette (M, yellow), and Padasterjochhaus (P, green).

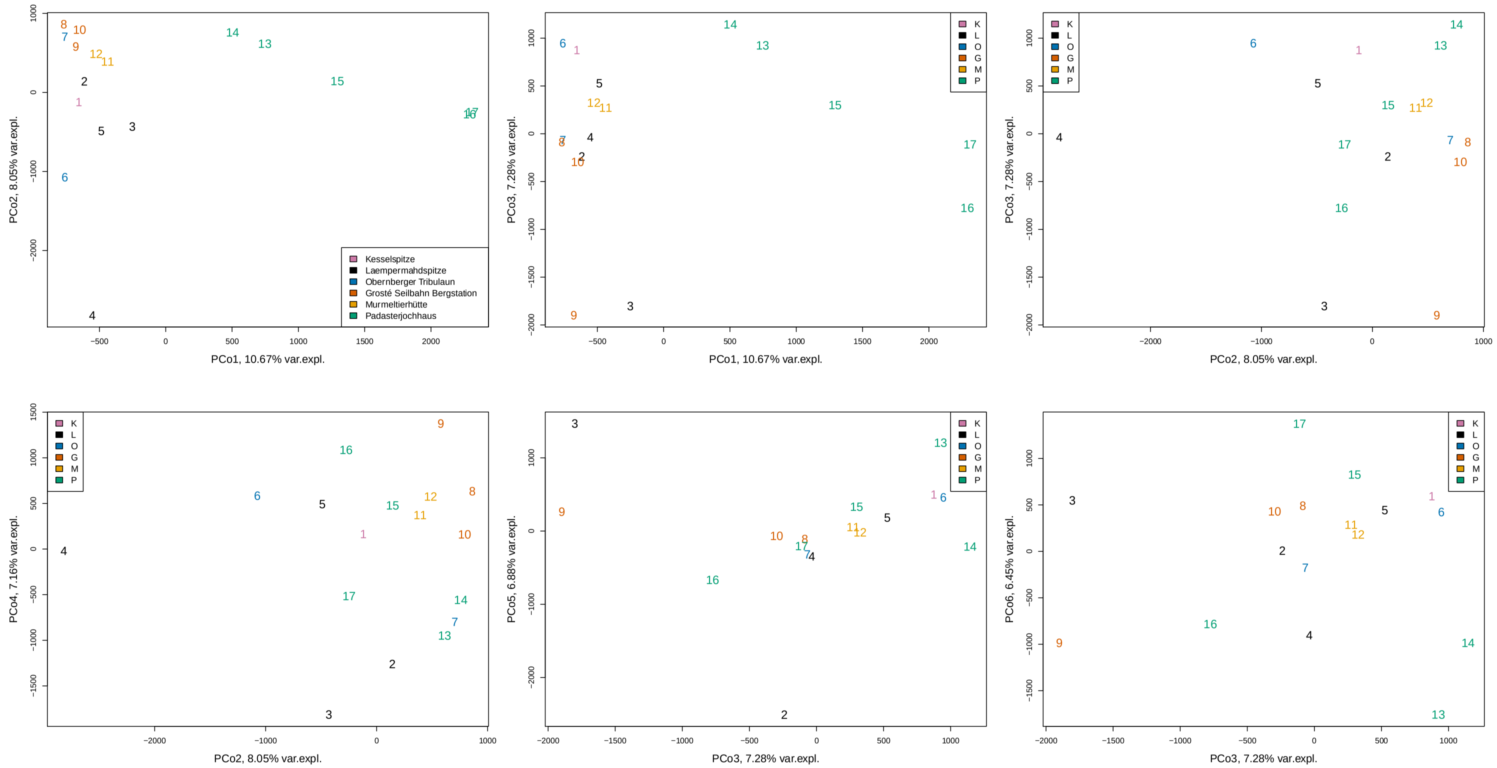


**Figure S3.** Principal coordinate analysis of 11,616 rare nuclear variants. Combinations of the first six principal coordinate axes are shown. Colors correspond to the different populations; Kesselspitze (K, pink), Laempermahdspitze (L, black), Obernberger Tribulaun (O, blue), Grosté Seilbahn Bergstation (G, orange), Murmeltierhuette (M, yellow), and Padasterjochhaus (P, green).

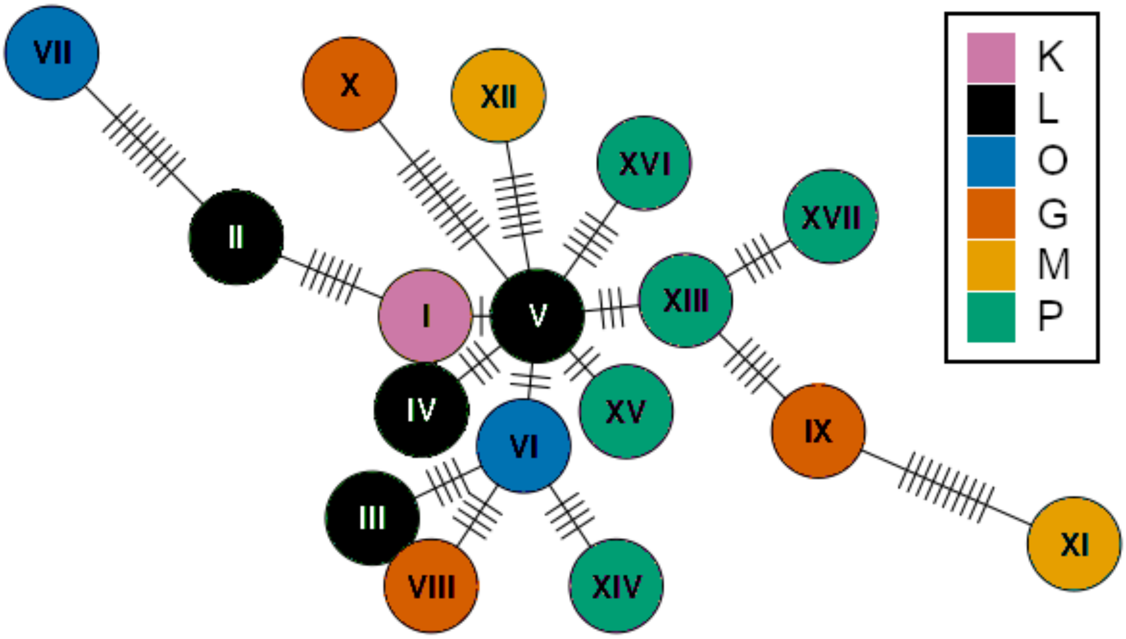


**Figure S4.** Haplotype network based on rare variants for 17 individuals and haplotypes Colors correspond to the different populations; Kesselspitze (K, pink), Laempermahdspitze (L, black), Obernberger Tribulaun (O, blue), Grosté Seilbahn Bergstation (G, orange), Murmeltierhuette (M, yellow), and Padasterjochhaus (P, green).

### 2. Supplementary Tables

**Table S1.** Information on sample IDs, sampling locations and read numbers during read processing.

| ID | Individual | Longitude | Latitude | Locality | Country | Elevation (m a.s.l.) | N reads raw (single) | N reads after quality filtering | N reads after decontamination | N reads after normalization |
|---|---|---|---|---|---|---|---|---|---|---|
| 1 | 92360 | 11.377325 | 47.102333 | K | AT | 2280 | 18356467 | 36247250 | 36247230 | 6629620 |
| 2 | 92361 | 11.378703 | 47.107025 | L | AT | 2220 | 26141248 | 51662040 | 51662022 | 9576382 |
| 3 | 92362 | 11.379536 | 47.108 | L | AT | 2220 | 25381606 | 50178814 | 50178794 | 9812710 |
| 4 | 92363 | 11.379769 | 47.10695 | L | AT | 2220 | 28471747 | 56414038 | 56414014 | 11374509 |
| 5 | 92364 | 11.380089 | 47.107153 | L | AT | 2220 | 18604788 | 36863302 | 36863290 | 8234937 |
| 6 | 92365 | 11.385581 | 46.987497 | O | AT | 2000 | 20013276 | 39559548 | 39559530 | 8164247 |
| 7 | 92366 | 11.385383 | 46.987583 | O | AT | 2000 | 23176562 | 45803428 | 45803404 | 9023848 |
| 8 | 92367 | 10.893125 | 46.219956 | G | IT | 2400 | 20324156 | 40180388 | 40180372 | 7418644 |
| 9 | 92368 | 10.889464 | 46.222336 | G | IT | 2400 | 25281428 | 49979044 | 49978924 | 9740077 |
| 10 | 92369 | 10.892083 | 46.220806 | G | IT | 2400 | 17042850 | 33707928 | 33707822 | 6535554 |
| 11 | 92370 | 11.700775 | 46.511847 | M | IT | 2200 | 16068604 | 31678722 | 31678652 | 5559225 |
| 12 | 92371 | 11.699558 | 46.511008 | M | IT | 2200 | 17989323 | 35660912 | 35660836 | 6805716 |
| 13 | 92372 | 11.358669 | 47.082906 | P | AT | 2320 | 13016877 | 25638612 | 25638592 | 5694169 |
| 14 | 92373 | 11.358606 | 47.083008 | P | AT | 2320 | 17648063 | 34718940 | 34718882 | 7340238 |
| 15 | 92374 | 11.358481 | 47.082889 | P | AT | 2320 | 16835713 | 33167428 | 33167402 | 7469014 |
| 16 | 92375 | 11.358458 | 47.082933 | P | AT | 2320 | 21532793 | 42572006 | 42571970 | 8901355 |
| 17 | 92376 | 11.358381 | 47.082917 | P | AT | 2320 | 26486877 | 52445844 | 52445818 | 10417939 |

**Table S2.** A summary of the annotation and gene ontology analysis can be found in Table S2 deposited at: https://github.com/mphaider/M.pallida.git

**Table S3.** Information on the location of the ten longest open reading frames (ORFs) on the *Machilis pallida* mitochondrium and corresponding best SmartBlast hits (LenNT=length in nucleotides, LenAA=length in amino acids).

| Label | Strand | Frame | Start | Stop | LenNT | LenAA | SmartBlast | Accession |
|---|---|---|---|---|---|---|---|---|
| ORF45 | - | 2 | 8392 | 6629 | 1764 | 587 | NADH dehydrogenase subunit V | YP_009047273.1 |
| ORF53 | - | 3 | 9807 | 8458 | 1350 | 449 | NADH dehydrogenase subunit IV | YP_009047274.1 |
| ORF25 | + | 3 | 10758 | 11894 | 1137 | 378 | cytochrome b | YP_009047277.1 |
| ORF13 | + | 2 | 554 | 1588 | 1035 | 344 | NADH dehydrogenase subunit II | YP_009047266.1 |
| ORF51 | - | 3 | 12948 | 11980 | 969 | 322 | NADH dehydrogenase subunit I | YP_009047278.1 |
| ORF14 | + | 2 | 3266 | 4120 | 855 | 284 | cytochrome c oxidase subunit II | YP_009047268.1 |
| ORF5 | + | 1 | 5059 | 5886 | 828 | 275 | cytochrome c oxidase subunit III | YP_009047271.1 |
| ORF15 | + | 2 | 4382 | 5059 | 678 | 225 | ATP synthase F0 subunit 6 | YP_009047270.1 |
| ORF20 | + | 3 | 2676 | 3332 | 657 | 218 | cytochrome c oxidase subunit I | YP_009047267.1 |
| ORF50 | - | 2 | 2410 | 1946 | 465 | 154 | cytochrome oxidase subunit I | SSC84599.1 |

Table S4: European Nucleotide Archive (ENA) accessions for samples and raw reads sequenced in this study.

| Primary accession | Secondary accession | Unique name | Study | Experiment |
|---|---|---|---|---|
| ERS4357532 | SAMEA6593248 | 92010 | ERP120116 | ERX4639242 |
| ERS4357531 | SAMEA6593247 | 92376 | ERP120116 | ERX4639621 |
| ERS4357530 | SAMEA6593246 | 92375 | ERP120116 | ERX4639620 |
| ERS4357529 | SAMEA6593245 | 92374 | ERP120116 | ERX4639619 |
| ERS4357528 | SAMEA6593244 | 92373 | ERP120116 | ERX4639618 |
| ERS4357527 | SAMEA6593243 | 92372 | ERP120116 | ERX4639617 |
| ERS4357526 | SAMEA6593242 | 92371 | ERP120116 | ERX4639616 |
| ERS4357525 | SAMEA6593241 | 92370 | ERP120116 | ERX4639615 |
| ERS4357524 | SAMEA6593240 | 92369 | ERP120116 | ERX4639614 |
| ERS4357523 | SAMEA6593239 | 92368 | ERP120116 | ERX4639613 |
| ERS4357522 | SAMEA6593238 | 92367 | ERP120116 | ERX4639612 |
| ERS4357521 | SAMEA6593237 | 92366 | ERP120116 | ERX4639611 |
| ERS4357520 | SAMEA6593236 | 92365 | ERP120116 | ERX4639610 |
| ERS4357519 | SAMEA6593235 | 92364 | ERP120116 | ERX4639609 |
| ERS4357518 | SAMEA6593234 | 92363 | ERP120116 | ERX4639606 |
| ERS4357517 | SAMEA6593233 | 92362 | ERP120116 | ERX4639600 |
| ERS4357516 | SAMEA6593232 | 92361 | ERP120116 | ERX4639597 |
| ERS4357515 | SAMEA6593231 | 92360 | ERP120116 | ERX4639565 |
| ERS5328265 | SAMEA7571643 | Combined_92360-92376 | ERP120116 | ERX4706073 |
